# Elucidating disease-associated mechanisms triggered by pollutants via the epigenetic landscape using large-scale ChIP-Seq data

**DOI:** 10.1101/2023.05.18.541391

**Authors:** Zhaonan Zou, Yuka Yoshimura, Yoshihiro Yamanishi, Shinya Oki

## Abstract

**Background:** Despite well-documented effects on human health, the action modes of environmental pollutants are incompletely understood. Although transcriptome-based approaches are widely used to predict associations between chemicals and disorders, the molecular cues regulating pollutant-derived gene expression changes remain unclear. Therefore, we developed a data-mining approach, termed “DAR-ChIPEA,” to identify transcription factors (TFs) playing pivotal roles in the action modes of pollutants.

**Methods:** Large-scale public ChIP-Seq data (human, *n* = 15,155; mouse, *n* = 13,156) were used to predict TFs that are enriched in the pollutant-induced differentially accessible genomic regions (DARs) obtained from epigenome analyses (ATAC-Seq). The resultant pollutant–TF matrices were then cross-referenced to a repository of TF–disorder associations to account for pollutant modes of action. We subsequently evaluated the performance of the proposed method using a chemical perturbation dataset to compare the outputs of the DAR-ChIPEA and our previously developed differentially expressed gene (DEG)-ChIPEA methods using pollutant-induced DEGs as input. We then adopted the proposed method to predict disease-associated mechanisms triggered by pollutants.

**Results:** The proposed approach outperformed other methods using the area under the receiver operating characteristic curve score. The mean score of the proposed DAR-ChIPEA was significantly higher than that of our previously described DEG-ChIPEA (0.7287 vs. 0.7060; *Q* = 5.278 × 10^−42^; two-tailed Wilcoxon rank-sum test). The proposed approach further predicted TF-driven modes of action upon pollutant exposure, indicating that (1) TFs regulating Th1/2 cell homeostasis are integral in the pathophysiology of tributyltin-induced allergic disorders; (2) fine particulates (PM_2.5_) inhibit the binding of C/EBPs, Rela, and Spi1 to the genome, thereby perturbing normal blood cell differentiation and leading to immune dysfunction; and (3) lead induces fatty liver by disrupting the normal regulation of lipid metabolism by altering hepatic circadian rhythms.

**Conclusion:** Highlighting genome-wide chromatin change upon pollutant exposure to elucidate the epigenetic landscape of pollutant responses outperformed our previously described method that focuses on gene-adjacent domains only. Our approach has the potential to reveal pivotal TFs that mediate deleterious effects of pollutants, thereby facilitating the development of strategies to mitigate damage from environmental pollution.

## BACKGROUND

Short- and/or long-term exposure to environmental pollutants has been associated with assorted adverse human health outcomes, including increased respiratory symptoms, heart or lung disease, and even early mortality [1–4]. Toxic pollutants in particular are implicated in carcinogenesis and other serious health effects, such as reproductive disorders and congenital anomalies [5–7]. To elucidate the modes of action (MoAs) by which chemicals, including environmental contaminants, elicit such toxic responses, computational strategies utilizing biomedical big data analysis have attracted the attention of researchers in recent decades [8–10]. For example, with the understanding that bioactive chemicals, such as pharmaceuticals and pollutants, affect human physiology by either maintaining or disrupting normal gene expression, respectively, comparative analyses of transcriptional profiles before and after chemical exposure, followed by gene ontology (GO) and pathway enrichment analyses, have been performed to elucidate chemical MoAs [11, 12]. However, although GO and pathway analyses can comprehensively extract functional characteristics from profiles of chemically induced differentially expressed genes (DEGs), considerable effort is required to identify specific primary literature that demonstrates the association between input DEGs and associated GOs or pathways. In addition, graph analytics (network analysis) is also frequently employed to estimate “molecular hubs” within chemically responsive gene regulatory networks, despite limitations in terms of the biological interpretability of the prediction outcomes [13].

Bioactive chemicals, including some pollutants, can target a wide variety of molecules, such as membrane transport proteins, ion channels, and metabolic enzymes. In particular, research on steroid receptors, which are activated by female (estrogens) and male (androgens) hormones, is actively pursued owing to the reported effects of endocrine-disrupting chemicals, also known as environmental hormones, on reproduction and development [14, 15]. In addition, heavy metals, including mercury and cadmium, which have been associated with pollution-related disorders such as Minamata disease and Itai-itai disease, may induce epigenetic changes as a component of their disease mechanisms [16, 17]. Furthermore, both intranuclear steroid receptors and enzymes that regulate epigenomic states, such as DNA methylation and histone modifications, can function as transcription factors (TFs) to directly control gene expression. Accordingly, certain pharmaceuticals such as estrogen receptor inhibitors and histone deacetylase inhibitors have been shown to exert anticancer effects by directly regulating TF function [18, 19]. Together, these observations suggest that TFs that collectively regulate the on/off switches of genes whose expression is significantly altered by exposure to environmental chemicals may function as direct targets or mediators of chemical MoAs.

To identify TFs whose binding is enriched to a given set of genes or genomic regions, binding motif-based methods (MotifEA) such as HOMER and MEME have widely been used [20, 21]. This approach predicts TF binding by extracting shared short nucleotide sequences, commonly referred to as “motifs”, from multiple input genomic regions. These motifs are then compared to known TF–DNA binding motifs by comparative analysis. As an alternative method for identifying key TFs in chemical MoAs, we recently developed an approach that is independent of TF–DNA binding motifs, which we termed the DEG-ChIPEA method [22]. This method is based on the ChIP-Atlas database, which extracts actual TF binding sites on the reference genome by curating and integrating tens of thousands of previously published ChIP-Seq experimental data examining TF binding to genomic DNA [23]. The DEG-ChIPEA method involves inputting chemically induced DEGs and performing enrichment analysis based on all ChIP-Seq experiments obtained from ChIP-Atlas (i.e., ChIP-Seq-based enrichment analysis; ChIPEA) to identify TFs whose binding is significantly enriched in the vicinity of the input DEGs. These identified TFs can be considered promising candidate pivotal factors that mediate the expression of therapeutic or adverse chemical effects. However, despite the generally high accuracy of DEG-ChIPEA, a potential limitation is that only TFs in the immediate vicinity of the transcription start site (transcription start site ± 5 kb) of the input chemically induced genes are analyzed. Thus, potential contributions to gene expression by TFs that bind to distal transcriptional regulatory regions, such as enhancers and topologically associating domain boundaries located far from the gene locus, are not taken into account.

In this study, we aimed to gain further insights into the MoAs of chemicals, particularly those of environmental pollutants, with regard to health effects. To achieve this, we propose a novel approach, termed “DAR-ChIPEA”, that considers TF binding events throughout the genome, including both proximal and distal transcriptional regulatory regions of genes, in contrast to the limited purview of the DEG-ChIPEA method. The DAR-ChIPEA method involves extracting differentially accessible regions (DARs), which are genomic regions with significantly different chromatin accessibility prior to and following chemical exposure, using data from environmental chemical perturbation epigenome experiments, such as ATAC-Seq datasets. These DARs are then used as input for ChIPEA to generate a chemical–TF association matrix, which allows the identification of TFs whose binding is significantly enriched in the input DARs. In addition, we constructed a chemical–TF–disorder triadic association by incorporating known TF–disease associations into the resulting chemical–TF matrix. Our approach outperformed methods based on TF–DNA binding motifs, as well as the previously developed DEG-ChIPEA method [22], which focuses only on TFs that bind in the vicinity of chemically induced DEGs.

## METHODS

### Chemical- or pollutant-perturbed transcriptome and epigenome data

To identify genomic regions with significant differences in chromatin accessibility (DARs) in response to chemical or pollutant exposure as determined through epigenome perturbation experiments, we used ATAC-Seq datasets retrieved from the DBKERO (*n* = 2,078; DRA006903–DRA006930) [24, 25] and TaRGET (*n* = 383; GSE146508) databases [24, 25]. In addition, RNA-Seq datasets corresponding to the ATAC-Seq datasets were made available in these databases and used to identify DEGs induced by chemical or pollutant exposure as input to the DEG-ChIPEA or DEG-MotifEA analyses (Additional file 1: Supplementary Fig. S1; DBKERO, *n* = 2,012, DRA006875–DRA006902; TaRGET, *n* = 468, GSE146508). DBKERO harbors perturbation data of 99 chemicals in human cell lines, whereas TaRGET curates *in-vivo* exposure data for seven pollutants in blood and liver samples from mice. All data used in this analysis were archived in the National Center for Biotechnology Information Sequence Read Archive (NCBI SRA) [26] and are associated with accession IDs as listed in Supplementary Tables S1–S4 (Additional file 2).

### Genome-wide TF binding and chromatin accessibility experimental data from ChIP-Atlas

Genome-wide TF binding site information was obtained from the ChIP-Atlas database [23], which comprehensively integrates nearly all publicly available ChIP-Seq data. Peak call data in BED format (MACS2 [27]; *Q* -value <1 × 10^−10^) were used in this study. Data from ChIP-Seq experiments examining binding of 1,796 and 866 TFs on the human (GRCh38/hg38; hereafter referred to as hg38) and mouse genomes (GRCm38/mm10; hereafter referred to as mm10), respectively, were used in this study (hg38, *n* = 15,155; mm10, *n* = 13,156). In addition, the information related to accessible chromatin used in this analysis to identify DARs induced by chemical or pollutant exposure was also obtained from ChIP-Atlas, which also provides peak call data for ATAC-Seq.

### TF–disease associations

Association data between TFs and diseases were obtained from DisGeNET (v 7.0) [28], which is a database of genetic and genomic information associated with human diseases. DisGeNET integrates this information from other expert-curated repositories and PubMed literature. Only manually curated data that were labeled with referenced PubMed IDs were retrieved, resulting in a total of 12,857 associations between TFs and diseases, involving 1,004 TFs and 3,205 diseases.

### Known chemical–disease associations as standard data

Manually curated chemical–disease association data were acquired from the Comparative Toxicogenomics Database (CTD) [29]. In total, we incorporated 28,821 chemical–disease associations involving 69 chemicals (DBKERO) and seven pollutants (TaRGET), and 2,150 diseases in this study.

### Identification of chemical- or pollutant-induced DEGs

After obtaining the SRA accession IDs of the chemical or pollutant perturbation RNA-Seq data from DBKERO and TaRGET, the raw sequencing data archived in NCBI SRA were downloaded and decoded into FASTQ format using the “fastq-dump” command from the SRA Toolkit (v3.0.0) [30]. Quality control was then performed using the “fastq_quality_filter” command from the FASTX-Toolkit (v0.0.13) [31]. Alignment to the reference genomes (DBKERO, hg38; TaRGET, mm10) was performed using HISAT2 (v2.1.0) [32], and expression quantification was performed using featureCounts (v2.0.3) [33]. All commands were run with default settings and parameters. DEGs were detected using the R Package “edgeR” (v3.40.2) [34]. Information on the statistical significance used for identification of DEGs is detailed in the Statistical Analyses section.

### Identification of chemical- or pollutant-induced DARs

Referring to the SRA accession IDs for the chemical- or pollutant-perturbed ATAC-Seq data obtained from DBKERO and TaRGET, the peak call data (in BED format) for the corresponding ATAC-Seq experiments were acquired from ChIP-Atlas. Peak calling was performed using MACS2 [27] in ChIP-Atlas; only peaks with a *Q* -value of <1 × 10^−10^ were considered as statistically significant accessible chromatin regions. Next, common open chromatin regions across all replicates for each chemical or pollutant perturbation condition were extracted using BEDtools (v2.23.0) [35]. Differential analysis was then performed to identify DARs induced by the chemicals or pollutants. Note that the term “differential analysis” in this context refers to the identification of genomic regions with no overlap between groups with and without chemical or pollutant exposure. In this study, DARs that exhibited relaxed or condensed chromatin upon chemical or pollutant exposure are referred to as “opened” or “closed” DARs, respectively.

### TF enrichment analysis fully incorporating public ChIP-Seq experiments (ChIPEA)

Details of the ChIPEA procedure using the application programming interface are provided online [22, 36]. To exploit the extensive ChIP-Seq data in ChIP-Atlas [23], we performed ChIPEA on profile TFs whose binding sites were enriched around genomic regions or genes of interest. In particular, using the genomic coordinates of opened and closed DARs, or the gene symbols of up- and down-regulated genes by a query chemical or pollutant as input, we first counted the number of intersects between the input data and peak-call data (MACS2; *Q* -value <1 × 10^−10^) of all TF-related experiments archived in the ChIP-Atlas database using the "intersectBed" command of BEDTools (v2.23.0) [35]. Note that 5 kb upstream and downstream of the transcription start site were added for each gene when chemically induced DEGs were queried. Enrichment scores (−log_10_*Q*) were then calculated using the two-tailed Fisher’s exact probability test to test whether the two datasets (opened and closed DARs, or up- and down-regulated genes) overlapped with the ChIP-Seq peak-call data with equivalent proportions before performing the Benjamini-Hochberg procedure for multiple testing correction; fold enrichment values (opened DARs/closed DARs) were concurrently obtained. If a chemical– or pollutant–TF association was indicated by multiple ChIP-Seq experiments, only the highest enrichment score was adopted. Parameters used in application programming interface-based ChIPEA were as follows: “genome=hg38 (DBKERO) or mm10 (TaRGET); antigenClass=TFs and others; cellClass=All cell types; threshold=100.” Sample codes for DEG-ChIPEA and DAR-ChIPEA are provided as indicated in the Availability of Data and Materials statement below.

### TF enrichment analysis focusing on TF–DNA binding motifs (MotifEA)

In this method, TF binding to chemical- or pollutant-induced DARs and DEGs was estimated using MotifEA with STREME and TOMTOM in MEME Suite (v5.4.1) [21], using default settings termed DAR-MotifEA and DEG-MotifEA, respectively. STREME was first run to identify 1,000 sequence motifs from queried multiple genomic regions, i.e., chemical- or pollutant-induced DARs or DEGs. Subsequently, TFs with enriched binding to the input genomic regions or genes were estimated by matching the identified motifs with known TF binding motifs obtained from HOCOMOCO v11 [37] using TOMTOM. Note that 5 kb upstream and downstream of the transcription start site were added for each gene when chemically induced DEGs were queried. Sample codes for DEG-MotifEA and DAR-MotifEA are provided as indicated in the Availability of Data and Materials statement below.

### Establishment of chemical– or pollutant–TF–disorder triadic associations

Using TFs enriched in chemical- or pollutant-induced DEGs and DARs as mediators, the TF– disease associations retrieved from DisGeNET [28] were integrated with the chemical– or pollutant–TF matrices obtained as outcomes of MotifEA or ChIPEA, thereby establishing chemical– or pollutant–TF–disorder triadic associations. The enrichment scores obtained via ChIPEA or MotifEA were continuously used as a quantitative measure to assess the degree of newly constructed chemical– or pollutant–disorder associations.

### Calculation of global area under the receiver operating characteristic curve (AUROC) and area under the precision recall curve (AUPR) scores

All predicted chemical– or pollutant–disorder associations were first reorganized into a single matrix containing enrichment scores with dimensions of *m* × *n*, where *m* and *n* represent the total number of chemical or pollutant perturbation conditions and the number of disorders, respectively (Additional file 1: Supplementary Fig. S1). Among various TFs linking a specific chemical–disease pair, we selectively adopted the most statistically significant TF, i.e., the one with the highest enrichment score. We then utilized the maximum enrichment score as a statistical measure to represent the association between this chemical and the disease. After referencing known chemical–disease associations from CTD [29], chemical-wise ROC (which plots the true positive rate against the false positive rate) and PR (which plots the precision [positive predictive value] against the recall [sensitivity]) curves were generated (R Package “ROCR” v1.0-11) [38]. The performance results were summarized into a global AUROC score, in which 1 represents perfect classification and 0.5 represents random classification, and an AUPR score, in which 1 represents perfect inference and the proportion of positive examples in the standard data corresponds to random inference.

### Mouse phenotype ontology enrichment analysis

GO enrichment analysis on opened DARs induced by pollutant exposure was performed using the Genomic Regions Enrichment of Annotations Tool (GREAT) (v4.0.4) [39]. Default parameters were used in this analysis; namely, “Species assembly: mm10; Association rule: Basal+extension: 5000 bp upstream, 1000 bp downstream, 1000000 bp max extension, curated regulatory domains included.” Ontologies categorized as “Mouse Phenotype” were then extracted. Parent category information for individual ontology terms was obtained with reference to the Mouse Genome Informatics database [40].

### Statistical Analyses

Statistical analysis was performed using R (v4.1.3). Unpaired two-tailed Wilcoxon rank-sum tests were used to compare AUROC and AUPR scores between two groups, Bonferroni correction was performed to presented adjusted *P*-values (*Q* -values). Fisher’s exact test was used for 2 × 2 contingency analysis, and the Benjamini-Hochberg procedure was used for multiple testing correction. Sample sizes and *Q* -values are shown in the figures, and a *Q* - value < 0.05 was considered statistically significant. NS denotes no significance. Chemical- and pollutant-induced DEGs were identified using the R package edgeR (v3.40.2), and a *P*-value <0.05 (DBKERO) or an adjusted *P*-value <0.1 (TaRGET) with an absolute value of log_2_ fold change >1 was recognized as statistically significant.

## RESULTS

### Study Design: Elucidation of Chemical MoAs Focusing on Chemically Reorganized Chromatin Accessibility

To expand upon our prior approach (DEG-ChIPEA) [22], here, we present a novel analytical approach to elucidate chemical MoAs focusing on chemically reorganized chromatin assembly. Specifically, we utilize chemically induced DARs as input to cover the full spectrum of TF binding to both proximal and distal transcription regulatory regions (e.g., promoters and enhancers; Fig. 1a). Our approach identifies TFs that integratively regulate chemical- or pollutant-induced DARs by comprehensively analyzing tens of thousands of publicly available ChIP-Seq data obtained from the ChIP-Atlas database (Fig. 1b) [23]. In addition, we integrated the predicted chemical–TF associations with disease-associated TF data obtained from the DisGeNET database [28]. The resulting chemical–TF–disorder triadic associations were then statistically validated using a known chemical–disease association dataset from CTD as standard data (Additional file 1: Supplementary Fig. S1) [29].

**Fig. 1.**
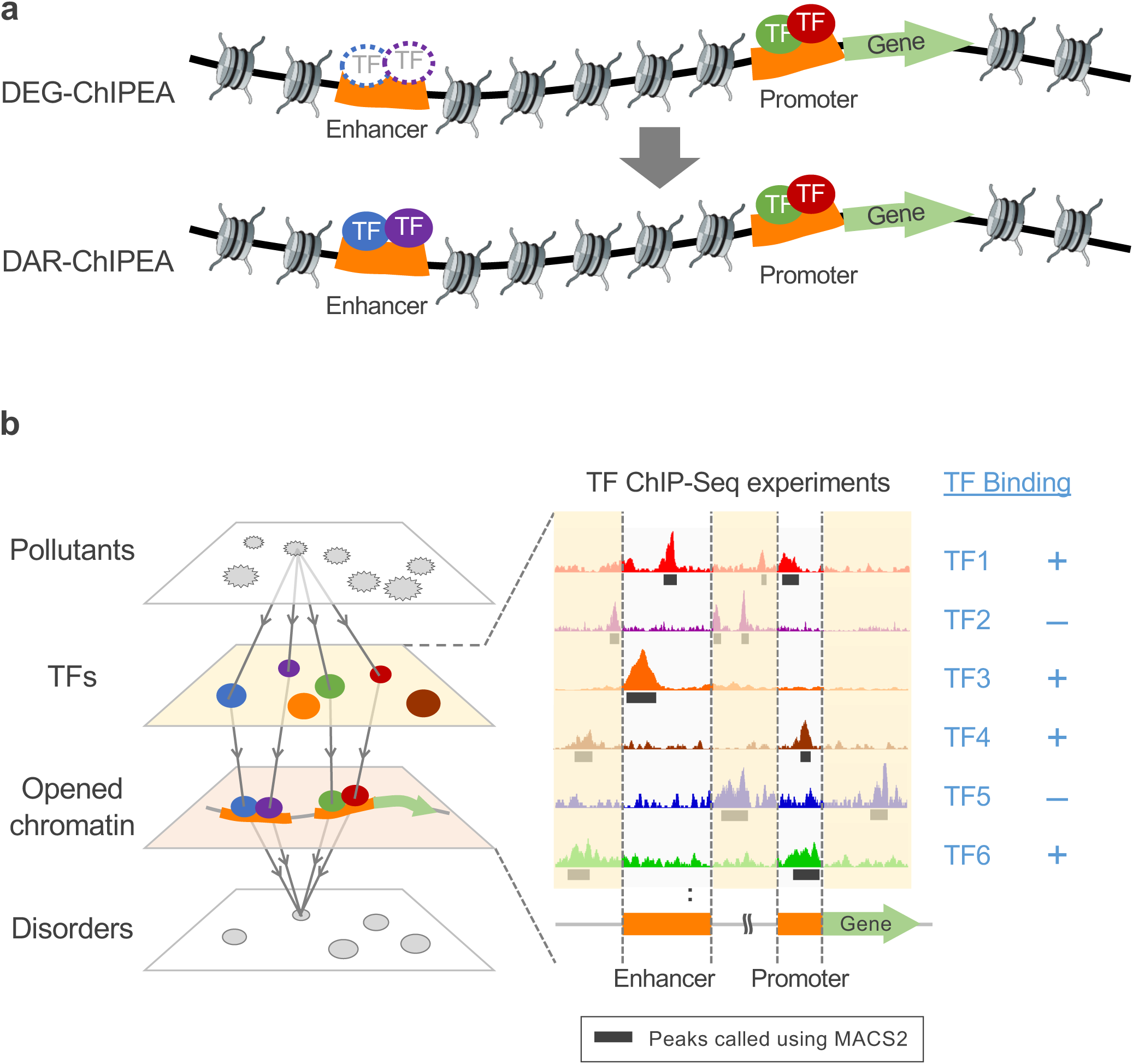
Overview of the proposed DAR-ChIPEA approach. **a** Schematic diagram illustrating the difference between our previously reported DEG-ChIPEA method [22] and the proposed DAR-ChIPEA approach. The DEG-ChIPEA method is restricted to analyzing TF binding to the vicinity of gene loci. In contrast, the DAR-ChIPEA approach comprehensively covers TF binding to accessible chromatin regions (obtained from ATAC-seq) across the genome including proximal (e.g., promoters) and distal (e.g., enhancers) regulatory elements. The circles surrounded by dotted lines represent the TFs binding to distal regulatory elements that are not detected in the DEG-ChIPEA method. **b** To demonstrate the feasibility of DAR-ChIPEA, we identified the TFs enriched at pollutant-induced DARs by analyzing large-scale ChIP-Seq data. Overlaps were evaluated between the DARs (green box) and peak-call data (black lines) of TF-related ChIP-Seq experiments archived in ChIP-Atlas

### Validation of the Proposed DAR-ChIPEA Approach to Predict TF-Driven Actions for ***In Vitro*** Chemical Perturbation

Because TaRGET [25] contains information on only a few types of pollutants, it is not sufficient to statistically evaluate the reliability of DAR-ChIPEA. To validate the proposed DAR-ChIPEA approach to predict TF-driven actions for *in vitro* chemical perturbation, we first evaluated the prediction accuracy of our method using the DBKERO database [24], which contains data on hundreds of chemical perturbations to cell cultures, providing correlated RNA-Seq and ATAC-Seq datasets. After retrieving pre-processed peak call datasets of ATAC-Seq from cell lines with and without chemical treatment in the DBKERO database from ChIP-Atlas, we performed a comparative analysis to detect DARs opened or closed by chemical perturbation (Additional file 1: Supplementary Fig. S2). Next, we fully exploited almost all publicly available human TF ChIP-Seq data in ChIP-Atlas to identify TFs enriched for binding to the extracted DARs using ChIPEA (DAR-ChIPEA). This allowed us to generate chemical–TF associations for each drug treatment condition. Furthermore, to evaluate the pivotality of TFs in drug MoAs, chemical–TF–disorder associations were constructed by assigning TF-related disease information obtained from DisGeNET to the resulting chemical–TF association matrix. For comparison, in the chemical–TF association generation step, we also used chemically induced DEGs as input to ChIPEA (DEG-ChIPEA), as previously reported (Additional file 1: Supplementary Fig. S2) [22]. For the same purpose, a more conventional and commonly used TF enrichment analysis method focusing on TF–DNA binding motifs (MotifEA) was also performed with DEGs or DARs as input (hereafter referred to as DEG-MotifEA and DAR-MotifEA; Fig. 2a).

**Fig. 2.**
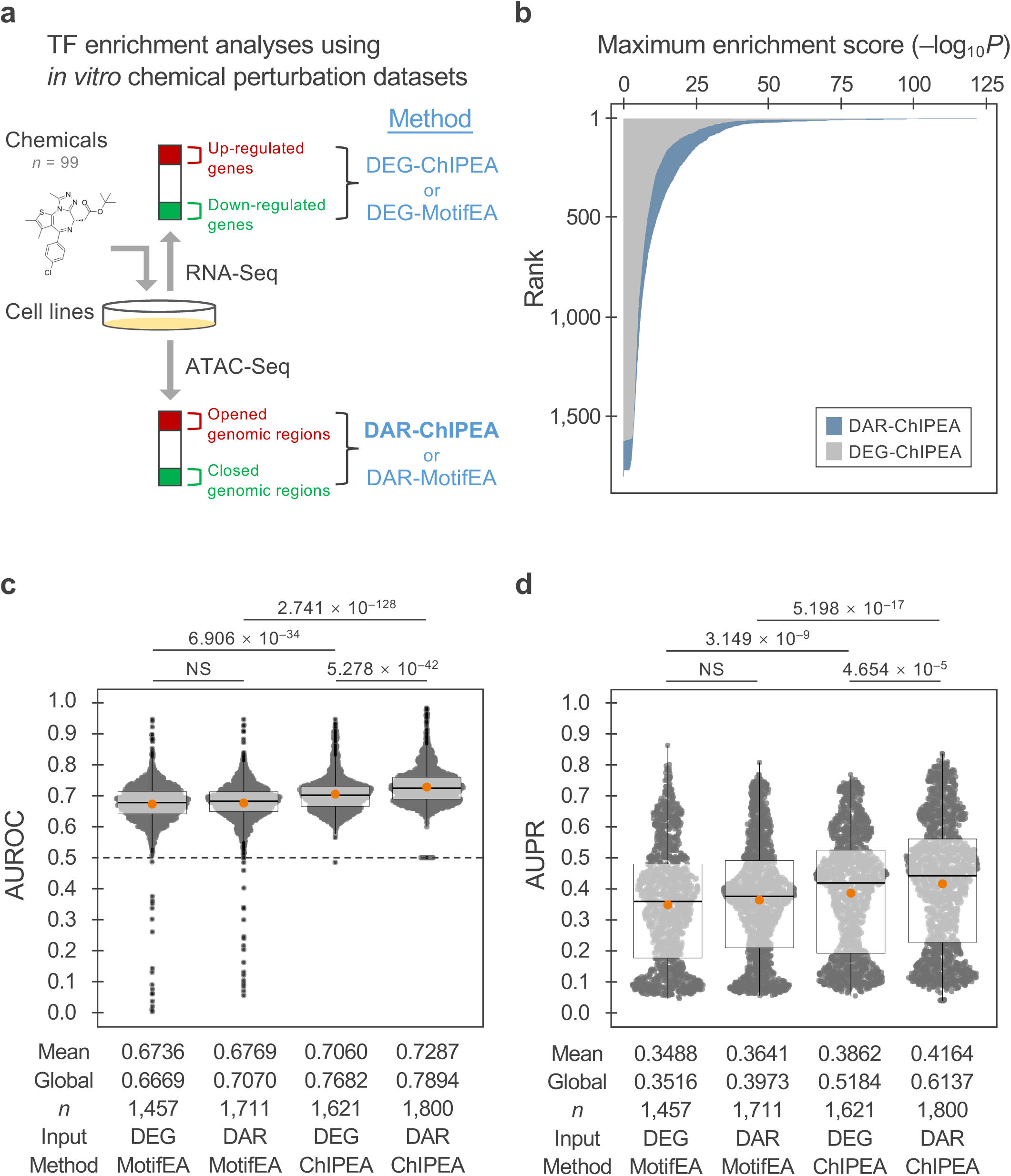
Validation of the proposed DAR-ChIPEA approach to predict TF-driven MoAs of *in vitro* chemical perturbation. **a** Schematic diagram for TF enrichment analyses using *in vitro* chemical perturbation datasets. DEGs or DARs upon chemical perturbation were detected using data from RNA-Seq or ATAC-Seq, respectively, followed by enrichment analyses to profile TF binding. Methods used for TF enrichment analysis are shown in blue; the proposed method (DAR-ChIPEA) is highlighted in bold. **b** Ranking plot of maximum enrichment scores (–log_10_*Q* -value) for each dosing condition obtained using the DAR-ChIPEA and DEG-ChIPEA methods. c and d Distribution of AUROC (**c**) and AUPR (**d**) scores for each chemical–TF– disorder triadic association predicted using DEG-MotifEA, DAR-MotifEA, DEG-ChIPEA, and ChIPEA. Orange dots represent mean scores; global scores are noted below the plots. Differences between the methods are presented as *Q* -values (two-tailed Wilcoxon rank-sum test) above the bee swarm plots

We observed that the statistically significant maximum enrichment scores (–log_10_*Q* -value) obtained from the proposed DAR-ChIPEA method were globally higher than those from our previously developed DEG-ChIPEA method (Fig. 2b; mean maximum enrichment score = 10.25 and 8.139, respectively; *P* = 3.013 × 10^−4^; two-tailed Wilcoxon rank-sum test), indicating that DAR-ChIPEA has a wider dynamic range than DEG-ChIPEA. Thus, DAR-ChIPEA offers a significant advantage in discriminating associated and non-associated TFs with regard to individual chemicals.

We then examined whether the chemical–TF–disorder triadic associations derived from DAR-ChIPEA that exhibited higher statistical significance were congruent with existing knowledge. The distribution of AUROC and AUPR scores for each dosing condition, generated using manually curated chemical–disease association data obtained from CTD as reference data (1,457, 1,711, 1,621, and 1,800 conditions for DEG-MotifEA, DAR-MotifEA, DEG-ChIPEA, and DAR-ChIPEA, respectively), was depicted using bee swarm plots (Fig. 2c, d; Additional file 2: Supplementary Table S5). The mean AUROC and AUPR scores of the DAR-ChIPEA approach across all dosage conditions were 0.7287 and 0.4164, respectively. These scores were significantly higher than those of other methods performed for comparison (DEG-MotifEA: mean AUROC 0.6736; mean AUPR 0.3488, DAR-MotifEA: mean AUROC 0.6769; mean AUPR 0.3641, and DEG-ChIPEA: mean AUROC 0.7060; mean AUPR 0.3862; *P*<0.05 for all; two-tailed Wilcoxon rank-sum test).

In addition, because the maximum enrichment scores varied drastically depending on the given dosing conditions, we were concerned that the discrimination of items was limited when comparing TFs within a dosing condition with low maximum enrichment scores. Therefore, we also calculated “global” statistics using an inter-chemical merged enrichment score vector to emphasize the significance of the actual values of the enrichment scores (detailed in the Methods section). The global AUROC and AUPR scores of the proposed approach were 0.7894 and 0.6137, respectively, significantly outperforming those of other methods (*P*<0.05; two-tailed Wilcoxon rank-sum test; Fig. 2c, d). These results suggest that the ChIPEA method using chemically induced DARs as input is a powerful approach for identifying pivotal TFs, elucidating key factors for specific chemical MoAs and predicting disorders or phenotypes associated with chemical treatments. The use of this approach may ultimately lead to increased predictive power compared to that of the existing DEG-ChIPEA and MotifEA-based computational methods.

### Application of the DAR-ChIPEA Approach to Predict TF-Driven MoAs upon Pollutant Administration *In Vivo*

We next employed the DAR-ChIPEA method to predict disorders caused by pollutants and TF-driven MoAs upon pollutant administration *in vivo* . After retrieving ATAC-Seq peak call datasets obtained with and without pollutant exposure from the TaRGET database available in ChIP-Atlas, we identified opened and closed DARs induced by various pollutants, as described in the Methods section (Additional file 1: Supplementary Fig. S2). We used publicly available mouse TF ChIP-Seq data in ChIP-Atlas to generate pollutant–TF associations using the proposed DAR-ChIPEA approach. We simultaneously performed analyses using the other methods for comparison, namely, DEG-ChIPEA, DEG-MotifEA, and DAR-MotifEA (Fig. 3a). The resulting pollutant–TF matrices were then used to generate pollutant–TF–disorder associations, focusing on pivotal TFs, with reference to TF–disease associations from the DisGeNET database. We evaluated the accuracy of the predicted pollutant–disorder associations using ROC and PR curves using known chemical-disease association datasets from CTD as a standard.

**Fig. 3.**
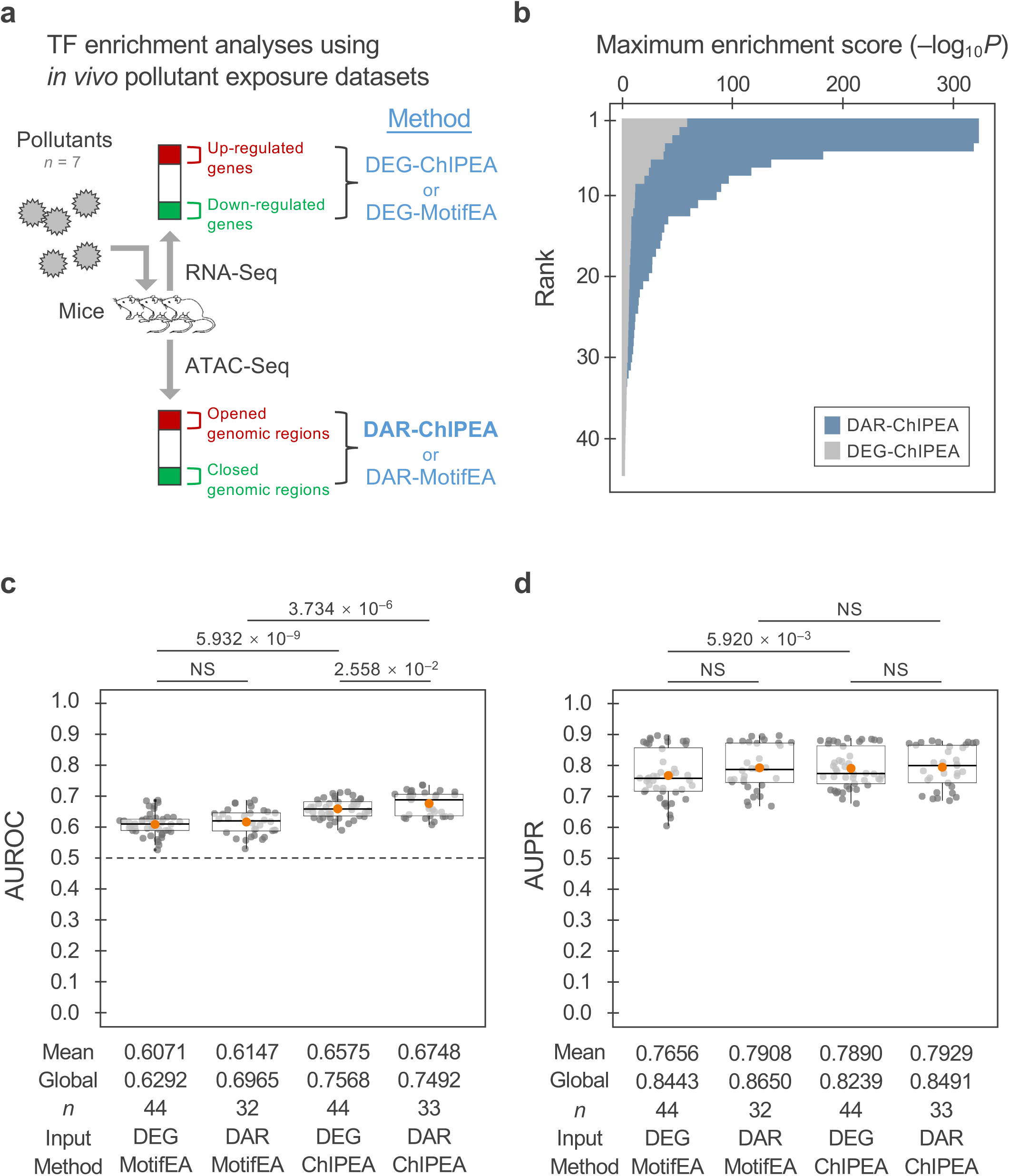
Application of the DAR-ChIPEA approach to predict TF-driven modalities of pollutant administration *in vivo* . **a** Schematic diagram for TF enrichment analyses using *in vivo* pollutant exposure datasets. DEGs or DARs upon pollutant exposure were detected using RNA-Seq or ATAC-Seq, respectively, followed by enrichment analyses to profile TF binding. Methods used for TF enrichment analysis are shown in blue, with the proposed DAR-ChIPEA method highlighted in bold. **b** Ranking plot of maximum enrichment scores (–log_10_*Q* -value) for each exposure condition as obtained using the DAR-ChIPEA and DEG-ChIPEA methods. c and d Distribution of AUROC (**c**) and AUPR (**d**) scores for each pollutant–TF–disorder triadic association predicted using DEG-MotifEA, DAR-MotifEA, DEG-ChIPEA, and ChIPEA. Orange dots represent mean scores; global scores are noted beneath the bee swarm plots. Differences between the methods are presented as *Q* -values (two-tailed Wilcoxon rank-sum test) above the bee swarm plots

The maximum enrichment scores attained using the proposed DAR-ChIPEA approach were globally higher than those achieved with our previously developed DEG-ChIPEA method, similar to the results obtained using *in-vitro* input data, further supporting the wider dynamic range for discrimination of associated TFs for individual pollutants (Fig. 3b; mean maximum enrichment score = 76.77 and 11.57, respectively; *P* = 8.421 × 10^−8^; two-tailed Wilcoxon rank-sum test).

In addition, we utilized a large set of publicly available Bisulfite-Seq experimental data (mm10; *n* = 34,793) integrated into ChIP-Atlas to assess the DNA methylation status within the DARs induced by representative pollutants. The results revealed that opened and closed DARs were enriched by either hyper- or hypo-methylated regions, suggesting DNA methylation change within pollutant-induced DARs (Additional file 1: Supplementary Fig. S3).

The distribution of AUROC and AUPR scores for each exposure condition (40, 28, 40, and 21 conditions for DEG-MotifEA, DAR-MotifEA, DEG-ChIPEA, and DAR-ChIPEA, respectively) was visualized using bee swarm plots (Fig. 3c, d; Additional file 2: Supplementary Table S6). The mean AUROC of the proposed DAR-ChIPEA approach across exposure conditions was 0.6748, significantly higher than those of the other methods (DEG-MotifEA, 0.6071; DAR-MotifEA, 0.6147; DEG-ChIPEA, 0.6575; *P*<0.05; two-tailed Wilcoxon rank-sum test). However, the DAR-ChIPEA AUPR score did not significantly (*P*>0.05) exceed those of the other methods (DEG-MotifEA, 0.7656; DAR-MotifEA, 0.7908; DEG-ChIPEA, 0.7890; proposed: DAR-ChIPEA, 0.7929). The global AUROC and global AUPR scores of DAR-ChIPEA were 0.7492 and 0.8491, respectively. These results suggest that the DAR-ChIPEA method performed satisfactorily in terms of AUROC; the absence of significant differences regarding AUPR may be attributed to the limited number of datasets analyzed.

### Biological Interpretation of the Predicted Pollutant–TF–Disorder Associations

In the proposed DAR-ChIPEA method, high AUROC scores were obtained for tributyltin (TBT) (blood samples), PM_2.5_ (blood samples), and lead (liver samples; AUROC = 0.7028, 0.7030, and 0.7349, respectively; Additional file 2: Supplementary Table S4, S6; highlighted with bold red font), which motivated us to evaluate the results of putative pivotal TFs involved in the MoAs of these pollutants in further detail, particularly with regard to the biological interpretation of the predicted pollutant–TF–disorder associations (Fig. 4; Additional file 2: Supplementary Tables S7–S9).

**Fig. 4.**
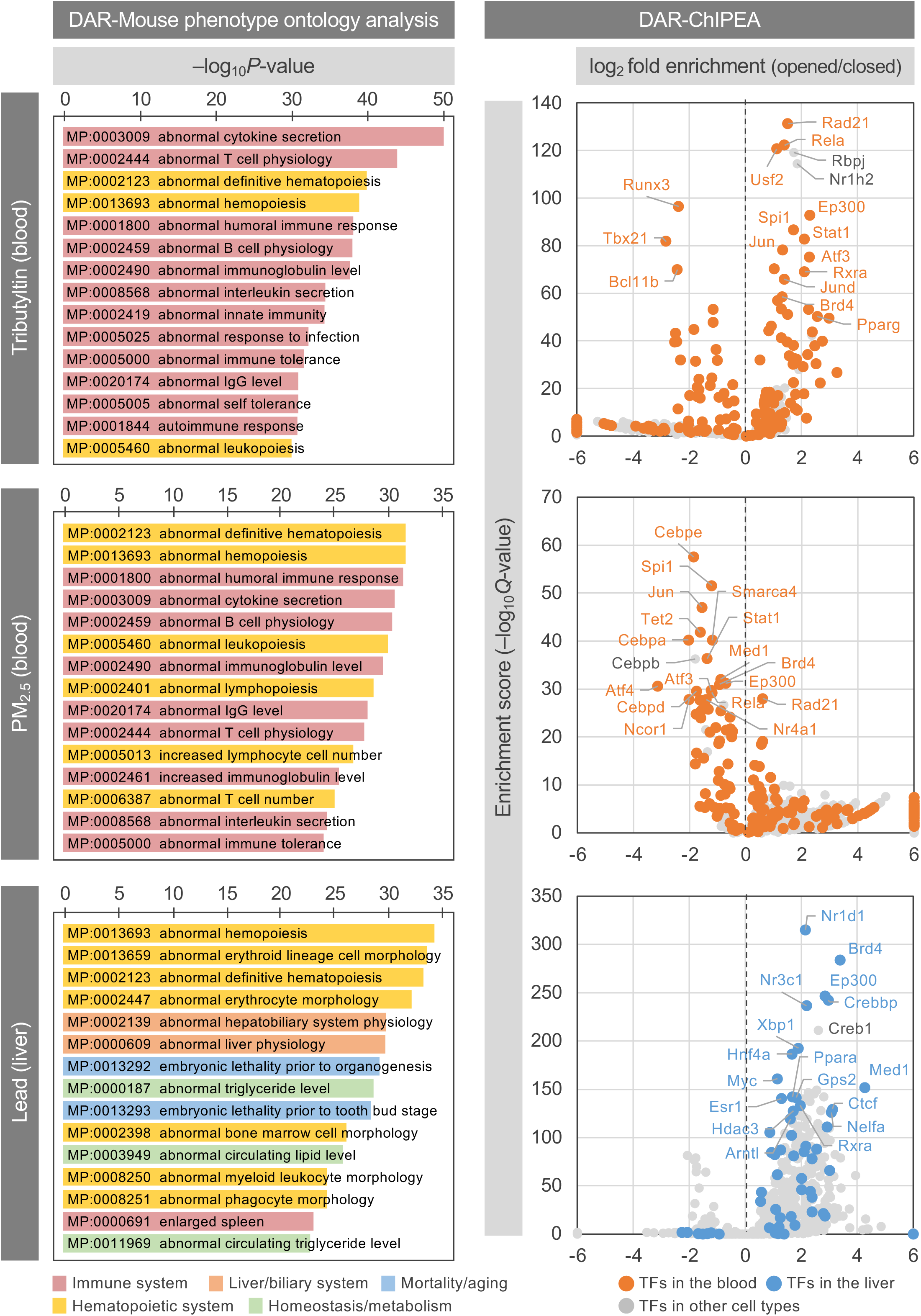
Biological interpretations of the predicted pollutant–TF–disorder associations for three representative pollutants. Mouse phenotype ontologies enriched for the genes proximal to pollutant-induced opened DARs are shown in the left panels. Bars representing individual ontology terms are colored based on their parent categories, which were obtained from the Mouse Genome Informatics database. Volcano plots illustrating the results of the proposed DAR-ChIPEA approach are shown in the right panels. Dots indicate individual TFs and are colored based on cell type classes obtained from ChIP-Atlas. Positive and negative values of log_2_ fold enrichment (opened DARs/closed DARs; X-axis) indicate TFs enriched in relaxed and condensed chromatin induced by pollutant exposure, respectively. The enrichment score (– log_10_*Q* -value; Y-axis) indicates the statistical significance of ChIPEA.

The organotin compound TBT is widely utilized in various industrial applications such as pesticides, paints, preservatives, and heat stabilizers for plastics. Nevertheless, TBT has been associated with a potentially increased risk of a number of lifestyle-related diseases, including obesity, fatty liver, cardiovascular disease, and abnormal glucose metabolism [41–44]. We initially conducted ontology enrichment analysis to discern any common nature of genes located proximal to opened DARs of blood samples upon TBT exposure using mammalian phenotype (MP) ontology. Notably, we found that the enriched gene sets were prominently manifested in phenotypic features of the immune system and hematopoietic system (upper left panel in Fig. 4). To further understand the transcriptional regulatory mechanisms involved in the response to TBT exposure, we then performed DAR-ChIPEA (upper right panel in Fig. 4). The genes encoding the top-20 enriched TFs exhibited robust expression in both control and TBT-exposed samples, and the same applied for PM_2.5_ and lead, as detailed below (Additional file 1: Supplementary Fig. S4). The results of DAR-ChIPEA showed that the opening of specific DARs susceptible to TBT exposure could enhance the enriched binding of Pparg, a master regulator of adipogenesis [45]. This is consistent with the *in-vitro* findings that TBT stimulates lipid accumulation by activating Pparg [46].

In addition, TBT can induce apoptosis in diverse tissues and cell categories, such as human peripheral blood lymphocytes and human amnion cells, and high-dose exposure to TBT has been reported to induce T lymphocyte apoptosis and thymic atrophy [47–49]. Furthermore, TBT suppresses the differentiation of naive CD4^+^ cells into Th1 cells, while stimulating differentiation into Th2 cells, thereby causing allergic disorders through the overproduction of Th2 cells, although the MoAs remain incompletely understood [50]. Our results showed that TBT exposure led to the enrichment of TFs such as Rela, Stat1, and Rbpj in opened DARs, consistent with the pro-inflammatory nature of TBT. Conversely, the binding of Bcl11b and Tbx21 was markedly enriched in closed DARs. BCL11B has apoptosis-inhibitory properties and is involved in maintaining T cell differentiation, as supported by the reduced numbers of T cells observed in *Bcl11b* -deficient mice [51, 52]. Notably, TBX21 also plays a key role in Th1 cell differentiation, with *Tbx21* -deficient naive Th cells exhibiting distorted Th2 cell polarization, a phenotype somewhat similar to that observed upon TBT exposure [53, 54]. These findings suggest that Bcl11b and Tbx21 may contribute to TBT-induced disruption of the Th1/Th2 balance.

PM_2.5_ is fine particulate matter suspended in the atmosphere with a diameter of 2.5 μm or less, which has the ability to transport pollutants in the environment. Exposure to PM_2.5_ is associated with respiratory diseases such as asthma, bronchitis, and pneumonia, as well as heart diseases such as arrhythmias and atherosclerosis, and significantly increases mortality from lung cancer [55–60]. Moreover, PM_2.5_ can worsen bacterial infections such as pneumonia caused by *Streptococcus pneumoniae* by impairing the function of macrophages in the lungs [61]. Oxidative and inflammatory damage and increased cellular autophagy and apoptosis have been implicated in the induction of various diseases by PM_2.5_; however, the underlying MoAs remain to be elucidated. Notably, the results of the MP ontology enrichment analysis for PM_2.5_ exhibited strong similarity to those of TBT, highlighting the prominent enrichment of phenotypes related to the immune and hematopoietic systems following PM_2.5_ exposure, consistent with previous research (middle left panel in Fig. 4) [62]. Furthermore, the results of DAR-ChIPEA revealed that the binding of blood cell differentiation-related factors such as Cebpe, Cebpb, Cebpa, Rela, and Spi1 was enriched in the DARs of blood samples closed by PM_2.5_ exposure, i.e., showed binding in the unexposed samples (middle right panel in Fig. 4). Among the enriched TFs, the C/EBP family members Cebpe and Cebpb are essential factors for differentiation and functional maintenance of granulocytes and the monocyte–macrophage axis, respectively, whereas Cebpa contributes to the regulation of growth arrest and differentiation of various cell types, including progenitor cells in bone marrow [63–65]. In addition, Rela is important to erythropoiesis, and Spi1 is important to early differentiation of B cells and bone marrow cells [66, 67].

Lead (Pb) has historically been used in a variety of applications, including paints and cosmetic dyes, and remains a common component of water pipes, solder, and other materials. Pb poisoning is typically a chronic condition that may not present with acute symptoms. Regardless of the absence of acute symptoms, Pb poisoning ultimately exhibits irreversible effects such as cognitive impairment, peripheral neuropathy, and progressive renal dysfunction [68–70]. Chronic Pb exposure may also induce fatty liver disease, which may be associated with changes in the gut microbiota [71]. Consistent with these observations, MP ontology enrichment analysis on liver samples exposed to Pb corroborated the significant enrichment of genes associated with the regulation of liver physiology, as well as lipid homeostasis and metabolism, in the proximity of DARs induced by Pb administration (lower left panel in Fig. 4). DAR-ChIPEA further revealed that Nr1d1 (also known as Rev-erbα), an important member of the feedback loop regulating the circadian clock composed of Clock and Arntl (also known as Bmal1) [72], was the most enriched TF in Pb-induced opened DARs. In addition, Ep300 and CREB-binding protein (Crebbp), transcriptional coactivators and members of the Clock/Bmal1 activator complex, were also found to be enriched in opened DARs (lower right panel in Fig. 4). These findings suggest that circadian rhythms in the liver may be disrupted by Pb exposure. Moreover, clock genes have been shown to act on the promoter and introns of adipogenic Ppara [73, 74]. In the liver, Ppara enhances lipid metabolism and is generally considered to have an inhibitory effect on hepatic steatosis, although some reports suggest that Ppara may promote the pathogenesis of non-alcoholic steatohepatitis [75, 76]. These findings, together with those from MP ontology analysis, suggest that Pb induces fatty liver not only through the gut microbiota pathway, but also by disrupting the normal regulation of lipid metabolism by circadian rhythms in the liver.

## DISCUSSION

In this study, we used the ATAC-Seq dataset, derived from samples perturbed by environmental chemicals, from public databases to identify pivotal TFs involved in chemical MoAs. We extracted DARs and applied ChIPEA, taking full advantage of large-scale ChIP-Seq experiments, to identify TFs that were enriched for binding to these genomic regions. In addition, we integrated the TF-associated disease dataset with the putative chemical–TF associations to estimate key TFs mediating chemical-induced disorder induction and in turn understand chemical MoAs. The proposed DAR-ChIPEA approach outperformed other existing computational methods in terms of accuracy.

In general, when cells or tissues are exposed to chemical substances, a series of events occur, initialized by pioneer factors, which are TFs that modulate chromatin accessibility. Subsequently, various cooperative TFs, often acting in complexes (e.g., Arntl/Ep300/Cebp in Fig. 4), bind to the newly opened or closed genome regions, causing alterations in the expression of genes regulated by these TFs. Moreover, the DNA methylation status undergoes modifications, contributing to the fixation of gene expression alteration. This entire cascade ultimately leads to phenotypic changes induced by chemical substances. The proposed method can provide insights into identifying key DNA-binding proteins involved in the MoAs of chemicals, including both pioneer factors and cooperative TFs. In addition, the public Bisulfite-Seq dataset in our previously developed ChIP-Atlas database can also be utilized to infer DNA methylation status change upon chemical exposure (Supplementary figure S2).

When dealing with health hazards caused by air pollution or industrial wastewater, it can be difficult to isolate a single pollution-causing molecule from such complex contexts. Application of approaches that focus on the detailed molecular structure of chemicals to analyze their MoAs, such as docking simulation and supervised learning using molecular structure as features, can thus be challenging [8–10, 77]. In this respect, the enrichment analysis-based approaches presented in this study are advantageous in that they use omics data as input and thus are better suited to analyze complex mixtures. In parallel with the processes used for causative molecule isolation, in the proposed DAR-ChIPEA method, chemically induced DAR information is obtained from experiments involving exposure to polluted air and water and passed through the proposed analysis pipeline, allowing for rapid and rough estimation of the key TFs involved in the MoAs of pollutants. This approach is expected to serve as an efficient complementary approach for investigating the causes of clinically-relevant pollution damage.

We found that the ChIPEA approach, an enrichment analysis method based on actual ChIP-Seq experiments, was more accurate than MotifEA, a numerical analysis method based on DNA binding motifs of TFs, in estimating chemical–disorder associations via key TFs. The moderate performance of MotifEA likely occurred, in part, because the presence of a particular binding motif does not necessarily indicate the actual binding of a TF, and conversely, it is not uncommon for a TF to bind to genomic regions that are distinct from known motifs. Furthermore, binding motifs can vary greatly depending on the tissue or cell type. In comparison, ChIPEA is highly reliable for studying the interaction between TFs and genomic DNA because it extracts TF binding sites in the genome based on a large number of ChIP-Seq experiments reported in the literature in a motif-independent manner [22, 36]. However, it should be acknowledged that the number of ChIP-Seq data currently available varies considerably across different tissues and cell types, and thus, ChIPEA may be less effective in situations in which available data are insufficient to generate statistically significant results.

Notably, the pivotal TFs found using our proposed approach may also serve as potential targets for drug discovery aimed at mitigating health hazards posed by environmental pollutants. Therefore, our approach is not only useful for elucidating the MoAs of environmental chemicals, but also for screening candidate compounds for potential pharmaceuticals. Owing to its ability to achieve an adequate depth of analysis with as few as 500 cells, ATAC-Seq provides the possibility to perform large-scale analyses of various compounds using small-well plates [78]. RNA-Seq may also be employed simultaneously to evaluate the pharmacological effects of the input compounds. In addition to the DBKERO database used in this study, other databases are also available that focus on predicting the medicinal effects and toxicity of various chemicals by analyzing changes in gene signatures upon chemical perturbation. For example, the LINCS L1000 project and Open TG-GATEs have obtained extensive transcriptome data on tens of thousands of drugs applied to various cell lines and laboratory animals [79, 80]. To further clarify the molecular cues that mediate gene expression changes in response to chemical administration, ATAC-Seq data should be incorporated into the compound screening pipeline in these projects, thereby facilitating the effective application of the proposed DAR-ChIPEA method. Furthermore, because ChIPEA is already publicly available through the ChIP-Atlas website in both a user-friendly graphical user interface and a command-based application programming interface, it could be easily applied in various situations to elucidate drug MoAs. Realization of this potential application could help identify TFs that are critical for the expression of drug effects, allowing the formulation of strategies to minimize unexpected risk, especially for compounds with potential adverse effects. In addition, identification of novel targets will allow the repositioning of drugs that have successfully passed safety testing as therapeutic agents for other diseases. Thus, the proposed DAR-ChIPEA approach could also contribute to lower drug discovery costs and a more efficient drug repositioning in the future.

## CONCLUSIONS

In this paper, we introduced a computational approach for highlighting genome-wide chromatin change upon pollutant exposure to elucidate the epigenetic landscape of pollutant responses, which outperformed other methods focusing on gene-adjacent domains only. The proposed method can be adopted to identify pivotal TFs involved in the MoAs of pollutants from an epigenetic perspective, thereby facilitating the development of strategies to mitigate environmental pollution damage. The proposed DAR-ChIPEA approach can also be used to further understand the MoAs of candidate drug compounds as well as to discover unexpected therapeutic and side effects of approved drugs to reduce costs during drug discovery.

## DECLARATIONS

### Ethics approval and consent to participate

Not applicable.

### Consent for publication

Not applicable.

### Availability of data and materials

We used freely available datasets; specific version numbers or dates for accessed or downloaded datasets are indicated as appropriate in the reference list. Sample codes and all raw results of the TF enrichment analyses (i.e., DEG-MotifEA, DAR-MotifEA, DEG-ChIPEA, and DAR-ChIPEA) were deposited at https://github.com/zouzhaonan/dar_moa/.

### Competing interests

The authors declare that they have no competing interests. The funders had no role in the design of the study, the collection, analysis, and interpretation of data, or in writing the manuscript.

### Funding

This work was supported by Grants-in-Aid for Scientific Research (KAKENHI), Japan Society for the Promotion of Science [22J15229 to ZZ, 23KF0048 to ZZ and SO, 22H02819, 23H04954 to SO]; Japan Science and Technology Agency [SPRING JPMJSP2110 to ZZ, PRESTO JPMJPR1942, NBDC JPMJND2202, and ERATO JPMJER2101 to SO]; Japan Agency for Medical Research and Development [PRIME JP22gm6710001, BINDS JP22ama121017 to SO], and Kyoto University [MIP and KMS-FUND to ZZ]. Funding for open access charge: Japan Science and Technology Agency [JPMJND2202].

### Authors’ contributions

Shinya Oki: Conceptualization, Supervision, Data curation, Funding acquisition, Writing– original draft, Writing–review & editing. Zhaonan Zou: Formal analysis, Funding acquisition, Writing–original draft, Writing–review & editing. Yuka Yoshimura: Formal analysis. Yoshihiro Yamanishi: Validation, Writing–original draft, Writing–review & editing.

## Acknowledgements

Computations were partially performed on the NIG supercomputer at ROIS National Institute of Genetics, Japan. We thank Prof. Shuh Narumiya and the members of the Department of Drug Discovery Medicine at Kyoto University for providing us with valuable advice.

## Supplementary Information

The online version contains supplementary material available at: **Additional file 1 (PDF): Supplementary Fig. S1** : Workflow of the proposed approach. First, we detected the pollutant-induced DARs using ATAC-Seq datasets obtained from the TaRGET database (*in vivo* pollutant exposure). We then performed ChIPEA with pollutant-induced DARs to predict TFs that play a pivotal role in the pharmacological effects of pollutants on health. The prediction phase was completed upon construction of pollutant– TF–disorder triadic associations integrating TF-disease associations retrieved from DisGeNET with the pollutant–TF matrices obtained as outcomes of ChIPEA. Subsequently, in the validation phase, we checked the authenticity of the predicted pollutant–disorder associations (shown as blue-colored continuous matrix) by referencing known chemical– disorder associations from CTD (shown as green-colored boolean matrix) and summarized the results as AUROC and AUPR scores. The *in vitro* chemical perturbation dataset obtained from the DBKERO database was also used to evaluate the prediction accuracy of the proposed method. The performance of other methods that utilize chemically induced DEGs as input or employ motif-based enrichment analysis (MotifEA) rather than ChIPEA were simultaneously assessed using the same workflow. **Supplementary Fig. S2** : Summary of the number of DEGs or DARs used as input to the enrichment analyses. Ranking plots are illustrated for individual chemical dosing or pollutant exposure conditions, and are sorted by the summation of the number of up- and down-regulated DEGs or opened and closed DARs. **Supplementary Fig. S3** : Results of the Bisulfite-Seq enrichment analysis for three representative pollutants. Dots indicate individual Bisulfite-Seq peak sets from ChIP-Atlas. Peak sets obtained from Bisulfite-Seq on blood (in the case of TBT and PM2.5 exposure) and liver (in the case of lead exposure) samples are colored (hyper-methylated regions, red; hypo-methylated, blue). Positive and negative values of log2 fold enrichment (opened DARs/closed DARs; X-axis) indicate peak sets enriched in opened and closed chromatin induced by pollutant exposure, respectively. The enrichment score (–log_10_*Q* -value; Y-axis) indicates the statistical significance of Bisulfite-Seq enrichment analysis. **Supplementary Fig. S4** : Expression levels of TFs mentioned in Figure 4 in mouse tissues before and after exposure to TBT, PM2.5, and lead. Genes encoding the top 20 enriched TFs in Figure 4 (right panels) are colored (rank 1–10, orange; rank 11–20, blue). CPM, count per million.

**Additional file 2 (XLSX): Supplementary Table S1:** *In vitro* chemical perturbation RNA-Seq dataset (DBKERO). **Supplementary Table S2:** *In vitro* chemical perturbation ATAC-Seq dataset (DBKERO). **Supplementary Table S3:** *In vivo* pollutant exposure RNA-Seq dataset (TaRGET). **Supplementary Table S4:** *In vivo* pollutant exposure ATAC-Seq dataset (TaRGET). Data shown in bold font are those that yielded high AUROC scores, as reported in the main text; see also Supplementary Table S6. **Supplementary Table S5:** AUROC and AUPR scores for *in vitro* chemical perturbation datasets (DBKERO). **Supplementary Table S6:** AUROC and AUPR scores for *in vivo* pollutant exposure datasets (TaRGET). Data shown in bold font are those that yielded high AUROC scores, as reported in the main text; see also Supplementary Table S4. **Supplementary Table S7:** Result of DAR-ChIPEA for exposing blood samples to TBT. **Supplementary Table S8:** Result of DAR-ChIPEA for exposing blood samples to PM_2.5_. **Supplementary Table S9:** Result of DAR-ChIPEA for exposing liver samples to lead.

## Notes

### Competing Interest Statement

The authors have declared no competing interest.

### Summary of Updates

Revised following reviewers' comments.

