## Supplementary figures for "Elucidating disease-associated mechanisms triggered by pollutants via the epigenetic landscape using large-scale ChIP-Seq data"

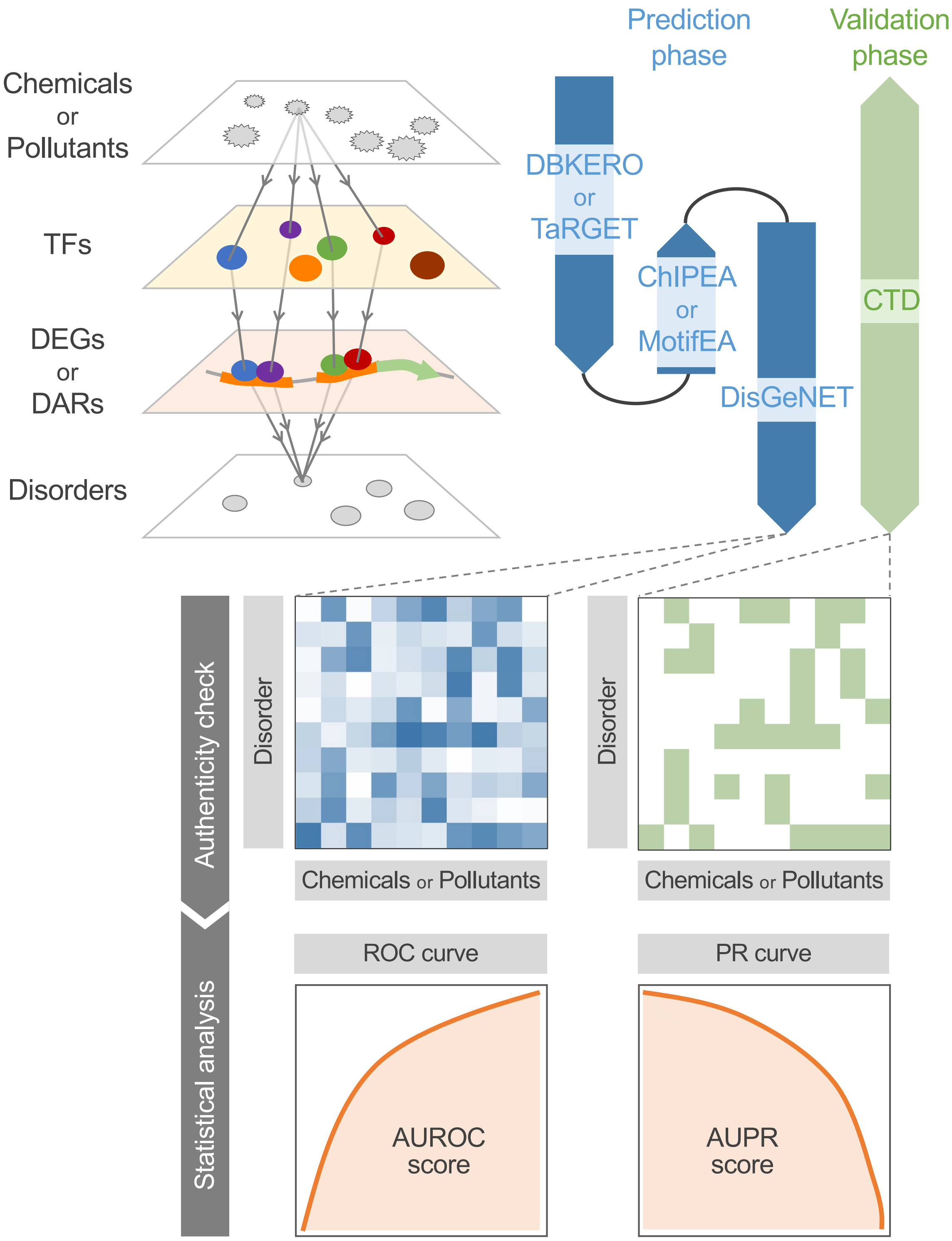


Supplementary Figure S1

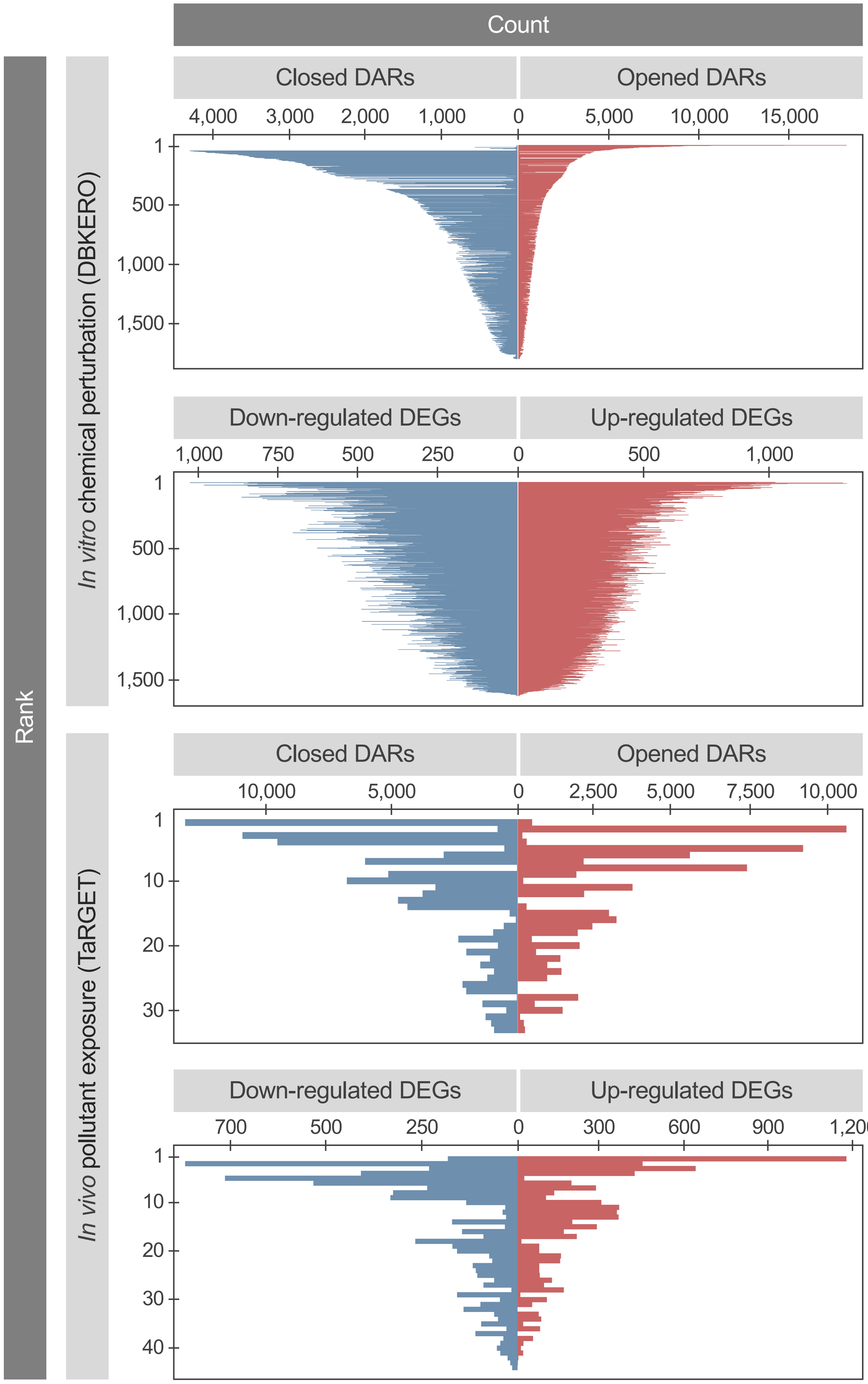


Supplementary Figure S2

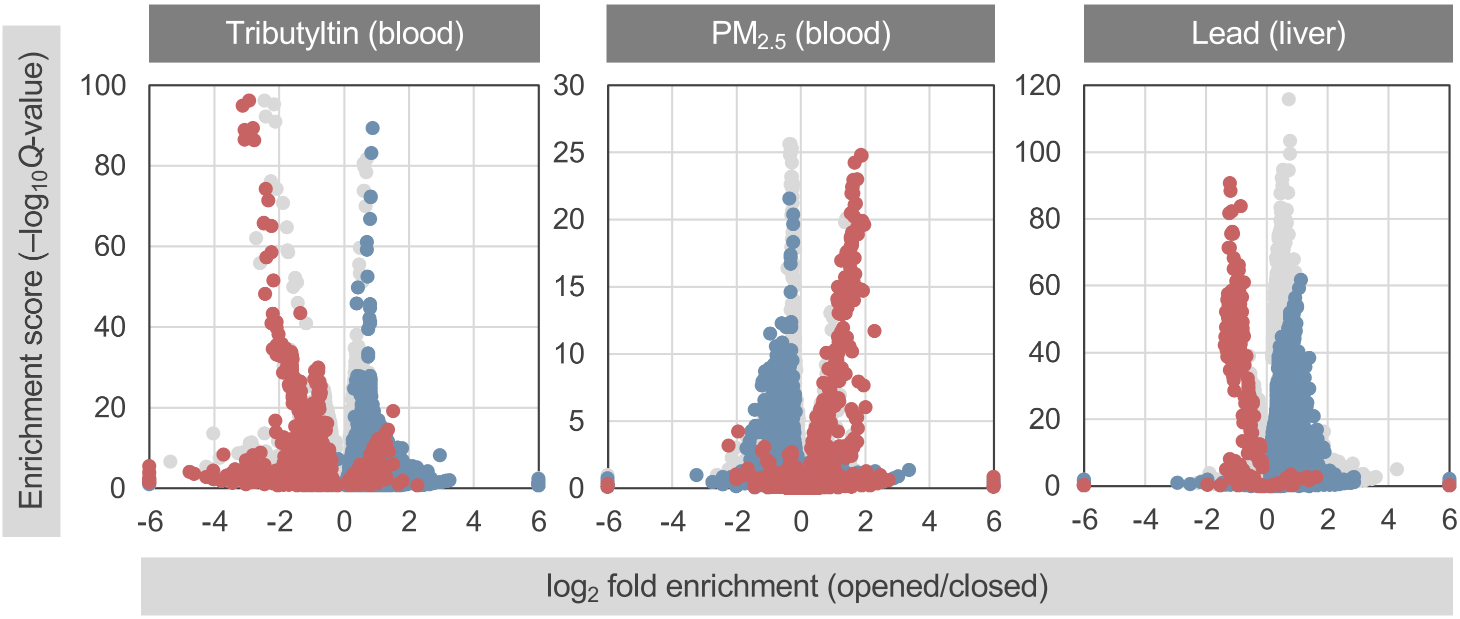


Supplementary Figure S3

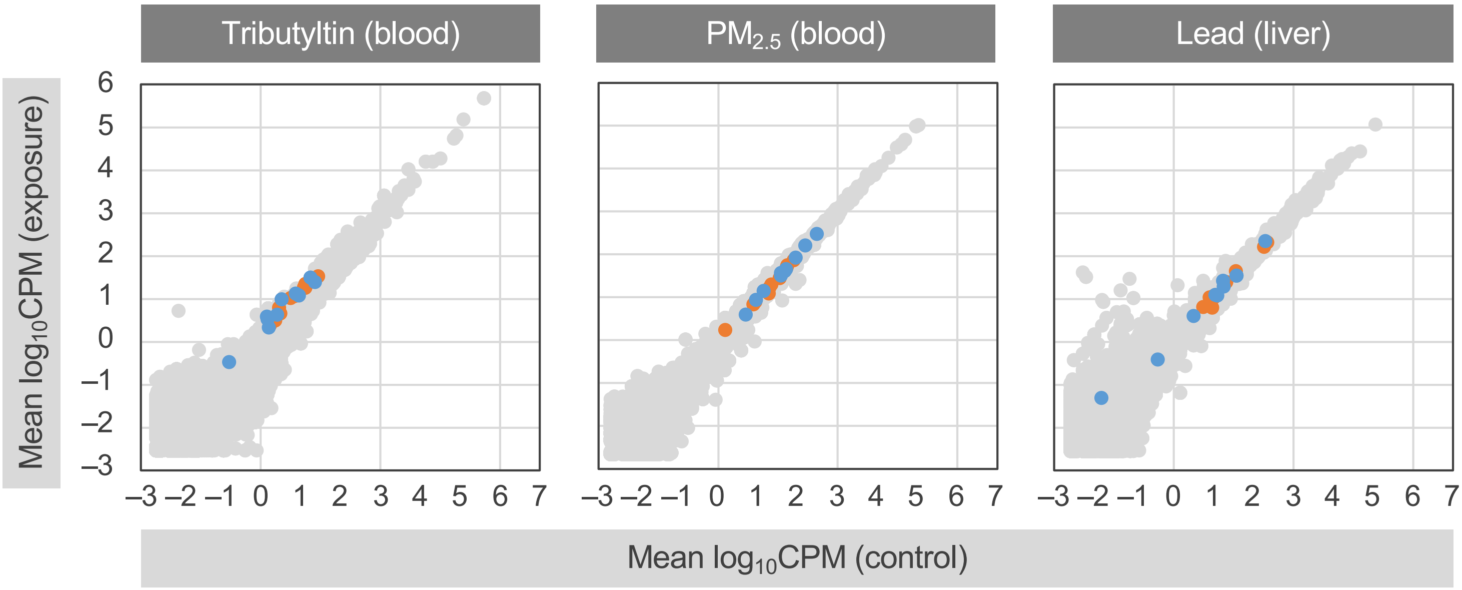


Supplementary Figure S4
